# Characterization of deglycosylated anti-cocaine mAb by differential scanning fluorimetry and fluorescence labeling

**DOI:** 10.1101/2025.10.21.683661

**Authors:** Terence L. Kirley

**Affiliations:** Department of Pharmacology, Physiology, and Neurobiology, College of Medicine, University of Cincinnati, 231 Albert Sabin Way, Cincinnati, OH 45267-0575

**Keywords:** Monoclonal antibody, deglycosylation, differential scanning fluorimetry, fluorescent labeling, antibody domains, cocaine binding

## Abstract

A humanized anti-cocaine mAb (h2E2) under development for the treatment of cocaine use disorders was deglycosylated enzymatically, and analyzed by differential scanning fluorimetry (DSF) using two distinct extrinsic dyes, and by FITC fluorescent labeling, to assess domain stability changes induced by deglycosylation of the Fc domain. The deglycosylated Fc fragment of the mAb was generated by proteolysis, and purified by ion exchange. Deglycosylation caused thermal destabilization of the glycosylated domain of the mAb, but there was no change in antigen (cocaine) binding or affinity. FITC fluorescent labeling of the deglycosylated mAb was suitable for qualitative assessment of cocaine binding. The thermal destabilization of the deglycosylated Fc fragment could be assessed by FITC labeling-based DSF, unlike the FITC-labeled control Fc fragment. This difference is due to the exposure of a secondary limited proteolysis site on the deglycosylated Fc, which increased accessibility and fluorescence labeling of Fc fragment lysine residues. Many of the methods and results in this study may be generalizable to numerous therapeutic mAbs, since the IgG1 anti-cocaine mAb shares the same glycosylation site and fully human heavy chain Fc sequence with other therapeutic human and humanized mAbs.

## 1. Introduction

We have generated and characterized a high (nM) affinity humanized anti-cocaine mAb named “h2E2”, intended for treatment of cocaine use disorders [1–3]. We also characterized the limited labeling of lysine residues of this anti-cocaine mAb by FITC [4]. This IgG1 mAb has a single glycosylation site on each heavy chain, which is conserved among members of this antibody class. We previously characterized the glycan chain structure and glycan structural heterogeneity of this anti-cocaine mAb by mass spectral analyses [5].

Glycosylation is known to be important for the solubility, tendency to aggregate, structural integrity, and thermal stability of many proteins, including antibodies [6–9]. Enzymatic removal of N-linked glycan chains using peptide N-glycosidase F (PNGase-F) is a commonly used, and very effective method of deglycosylation, which removes the entire, intact glycan chain, leaving an aspartic acid residue on the protein at the position of the originally glycosylated asparagine residue [10]. The effect of partial and full deglycosylation of an unrelated monoclonal antibody using a variety of biophysical techniques, such as circular dichroism, light scattering, fluorescence spectroscopy, and differential scanning fluorimetry has been previously reported [11].

In the current study, the effects of deglycosylation of the anti-cocaine mAb by peptide N-glycosidase F (PNGase-F) removal of the N-linked glycan chains was assessed, both on the intact mAb, and on the Fc mAb fragment containing the glycosylation sites. The deglycosylated mAb and Fc fragments were characterized by reducing and non-reducing SDS-PAGE, both before and after limited reaction with fluorescein isothiocyanate (FITC). The effect of deglycosylation on the mAb and the Fc fragment thermal stability and ability to bind the antigen, cocaine, was also assessed by differential scanning fluorimetry (DSF), using two distinct extrinsic DSF dyes, Sypro orange, and DASPMI, as well as by the covalently incorporated fluorescein fluorescence in the absence of any added extrinsic DSF fluorescent dye probes. Some of the methods and results reported in this study may be generalizable to numerous IgG1 mAbs, since the IgG_1_ anti-cocaine h2E2 mAb used in this study shares the same conserved glycosylation site and fully human heavy chain Fc sequence with many other fully human and humanized mAbs used therapeutically.

## 2. Experimental Procedures

### 2.1 Materials

The h2E2 anti-cocaine monoclonal antibody was supplied by the manufacturer, Catalent, as previously described [3], and dialyzed against 20 mM MOPS pH 7.4 buffer before use. Peptide N-glycosidase F (PNGase-F) was purchased from both New England BioLabs (Catalog P07042) and PROzyme (catalog GKE-5006B). Endo-Lys-C immobilized on agarose was purchased from SignalChem (catalog L585-31AN-1). DTT and all electrophoresis chemicals were from BioRad. The SDS-PAGE gel protein sizing standards were from SMOBIO (catalog PM1600). 10 mM stock solutions of cocaine in distilled water were made from solid cocaine-HCl (obtained from the Research Triangle Institute) as described [1]. Microcon 500 µl YM100 100 kDa molecular weight cutoff spin filters were from Amicon (catalog 42412), and the 0.22 µm 500 µl spin filters were Ultrafree MC with GV Durapore membranes (catalog UFC30GV0S) were obtained from Millipore. Zeba Spin 0.5 ml 7 kDa MWCO size exclusion desalting columns (catalog 89882) were from ThermoFisher Scientific.

Fluorescein isothiocyanate (FITC) was purchased from Invitrogen (isomer 1, catalog F1907). MOPS buffer was from ThermoFisher Scientific, and PBS buffer concentrate (10X) was purchased from Cambrex (BioWhittaker, without calcium or magnesium, catalog 17-517Q). Macro-Prep High S strong cation exchange resin (catalog 1560030) was from BioRad. Sypro orange 5000X dye stock was from Invitrogen/Life technologies, and the solid 4-(4- (dimethylamino)styryl)-N-methylpyridinium iodide dye (4-Di-ASP iodide, DASPMI) was purchased from ThermoFisher Scientific (Invitrogen/Life Technologies catalog D-288). DASPMI 20 mM stock dye solution was made by dissolution of the solid dye in dry DMSO, and Sypro orange and DASPMI dye stock DMSO solutions were stored light protected at −20°C. Applied Biosystems 48 well RT PCR plates (catalog 4375816) and the MicroAmp optical adhesive plate sealing transparent film (catalog 4375928) used for the StepOne RT PCR employed to perform the DSF analyses were purchased from ThermoFisher Scientific.

### 2.2 Methods

#### 2.2.1 mAb and Fab fragment protein concentrations and FITC degree of labeling (DOL) determinations

The concentration of the intact anti-cocaine h2E2 mAb was determined by absorbance at 280 nm, using an extinction coefficient of 219,500 M^-1^cm^-1^, while the concentration of Fc fragment was determined by absorbance at 280 nm using an extinction coefficient of 71,570 M^-1^cm^-1^. The degree of labeling (DOL, mol FITC/mol protein) of mAb or Fc with FITC was determined using these protein extinction coefficients, along with an extinction coefficient for fluorescein of 70,000 M^-1^cm^-1^ at 494 nm, and a correction factor of 0.3 for FITC at 280 nm (i.e., an extinction of 21,000 M^-1^cm^-1^ at 280 nm for fluorescein covalently attached to the mAb). Absorbance measurements were made using either 2 µL samples in an Implen N60 NanoPhotometer, or 50 to 100 µl samples in a 100 µl cuvette in a Beckman DU800 spectrophotometer.

#### 2.2.2 PNGase-F deglycosylation of anti-cocaine mAb

Deglycosylation of the intact mAb was carried out using PNGase-F from NEB or PROzyme on 1.0 ml of 14 mg/ml mAb in 20 mM MOPS 7.4 buffer for 16 hours at 22°C. The deglycosylated samples were then concentrated and separated from PNGase-F using YM100 100 kDa molecular weight cutoff spin filters, washed with 20 mM MOPS 7.4 buffer, and re-quantitated for mAb concentration by absorbance at 280 nm, with a measured final recovery of 75-80% of the starting mAb.

#### 2.2.3 SDS-PAGE analysis of mAb and Fc fragment samples

1-4 µg of each mAb and Fc fragment sample were analyzed on SDS-PAGE gels containing varying percentage concentrations of acrylamide, according to Laemmli [10]. Protein samples were diluted to 0.2 mg/ml and boiled for 3 minutes in either reducing (+ 100 mM DTT) or non-reducing (no DTT) sample buffer. The gels with FITC labeled proteins were photographed for fluorescence using the SYBR Green settings (excitation at 472 nm and emission at 595 nm) on an Azure 280 gel imager. All gels were stained for 1 hour with Coomassie blue, and destained overnight as described by Laemmli [13], prior to photography.

#### 2.2.4 Proteolysis with Endo-Lys-C and ion exchange to generate and purify the Fc fragment

Intact control and deglycosylated mAb at 2.0 mg/ml were treated with agarose bead-bound Endo-Lys-C protease for 60 minutes at 22°C in 20 mM MOPS pH 7.4. The fragments were then separated from the proteolytic enzyme beads by centrifugation through a 0.22 µm spin filter (GV Durapore). The resultant mixtures of Fab and Fc fragments were then mixed with Macro-Prep High S strong cation exchange resin (previously washed extensively with 20 mM Mops, pH 7.4) for 10 minutes with constant gentle mixing, and the resin removed by 0.22 µm spin filtration.

The unbound filtrate, containing the Fc fragment (the Fab fragment should be bound to the ion exchange resin under these conditions) was then re-incubated with a fresh aliquot of this same ion exchange resin to remove residual contaminating Fab and intact mAb. After resin removal as before, the purified Fc fragment was quantitated by A280 nm absorbance, and analyzed by SDS-PAGE under reduced and non-reduced conditions.

#### 2.2.5 FITC labeling of control and deglycosylated mAb and Fc fragments

Deglycosylated and control samples were labeled with FITC as follows. For the intact mAb, 100 µl of 5 mg/ml mAb in 100 mM NaBO_3_ pH 8.0 buffer was incubated with a final concentration of 2 mM FITC (from a freshly made 100 mM stock of FITC in DMSO) for 30 or 120 minutes at 22°C. The reaction was stopped by removing the FITC via size exclusion gel on 500 µl Zeba spin filters (7 kDa molecular weight exclusion), equilibrated with PBS buffer. The concentration of mAb and the stoichiometry of FITC labeling (degree of labeling, DOL) were determined using the extinction coefficients listed above (see section 2.2.1).

The purified Fc fragments (at a final concentration of 5 µM) were labeled with 100 µM FITC in either 20 mM MOPS pH 7.4 for 1 hour at 22°C, or to achieve a higher level of FITC labeling, in 100 mM NaBO_3_ pH 8.0 for 2 hours at 22°C. The Fc samples were then separated from excess FITC, exchanged into PBS buffer, and quantitated as described above for the intact mAb labeled with FITC.

#### 2.2.6 Differential scanning fluorimetry (DSF) of control and deglycosylated mAbs and Fc fragments

Differential scanning fluorimetry (DSF) was done using a ThermoFisher Scientific/Life Technologies StepOne RT PCR. When using the Sypro and DASPMI extrinsic dyes, the ROX dye calibration settings (fluorescence excitation at 485 nm, with emission at 610 nm) were used, and the passive reference was designated as none [4]. When using FITC labeled mAbs and Fc fragments, the FAM dye calibration settings (excitation at 470 nm, emission at 520 nm) were used, also with no passive reference [4]. In some experiments, cocaine was added to the samples, at final concentrations ranging from 3 to 1000 µM. For every sample analyzed, duplicate or quadruplicate 20 µl aliquots in pH=7.4 PBS buffer were loaded onto the 48 well plate. Either 2.5 µM or 1.0 µM mAb or Fc fragment in pH=7.4 PBS buffer was used, with the samples that were not labeled with FITC containing either 100 µM DASPMI dye, or a 1:500 dilution of commercial (5000X) Sypro orange dye stock. The DSF sample plates were heated from either 35°C to 95°C (for the intact mAb) or from 25°C to 95°C (for the Fc fragments), both at a heating rate of 0.45°C/minute, (machine heating ramp rate setting of 0.3%) as previously described [12].

## 3. Results

Control mAb and deglycosylated samples generated by two different commercial PNGase-F enzymes were analyzed using reducing SDS-PAGE, as shown in Figure 1. Different amounts of mAb proteins were loaded per lane to allow better visual assessment of the small shift in size caused by deglycosylation of the single glycan chain bound to each mAb heavy chain (HC), and therefore, the completeness of the deglycosylation process. The results indicated that the deglycosylation of the HC was complete using both commercial enzymes, with no change observed in the light chain, as expected. These samples were then analyzed by DSF using the Sypro orange dye, and titrating with cocaine, as shown in Figure 2. Several overlapping thermal unfolding processes detected with this dye are observed in the absence of cocaine in all samples, and these transitions are more completely separated from one another by adding cocaine, which thermally stabilizes a portion of the Fab fragment, moving its thermal unfolding transition to a higher temperature with increasing cocaine concentrations. This effect can be more clearly seen by just comparing the 0 and 30 µM cocaine results for the control and deglycosylated mAb (Figure 3). With 30 µM cocaine (Figure 3B), the lower Tm unfolding transition derivative peak shifts from approximately 68°C in the glycosylated mAb to 62°C in the deglycosylated mAb, indicating a large destabilization of this protein folding domain by removal of the glycan chain. There is no change in the Tm for the Fab domain derivative peak stabilized by cocaine binding (at approximately 76°C in the presence of 30 µM cocaine), suggesting no effect of deglycosylation on antigen (cocaine) binding.

**Figure 1.**
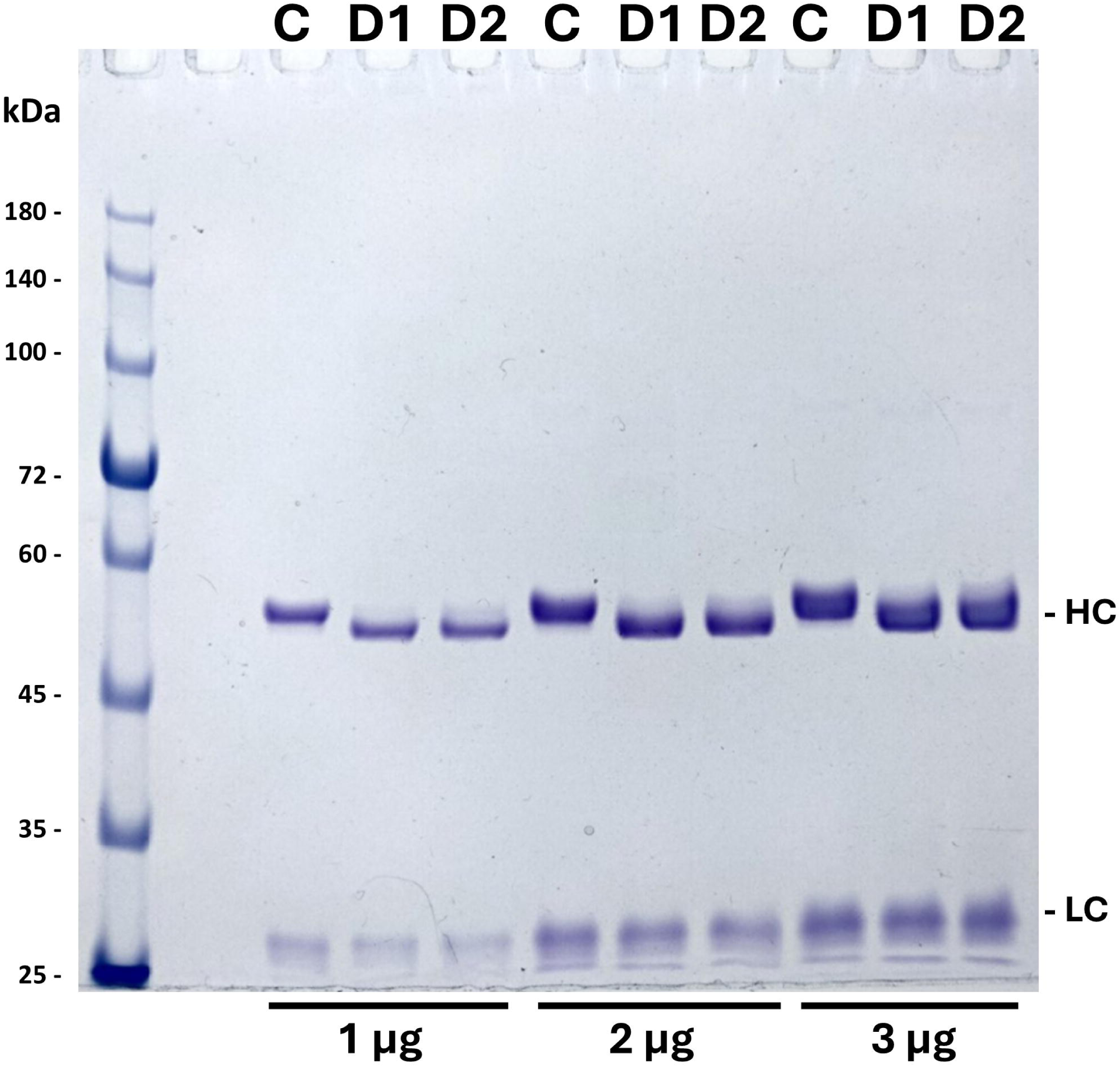
7% SDS-PAGE analysis of deglycosylated anti-cocaine mAb under reducing conditions. Analysis of 1, 2, or 3 ug of control (C) mAb or deglycosylated mAb (D1, deglycosylated with NEB PNGase F, D2, deglycosylated with PROzyme PNGase-F) as described in methods. Heavy chain (HC) and light chain (LC) bands are annotated. Note the shift to lower molecular weight for the HC in the deglycosylated samples.

**Figure 2.**
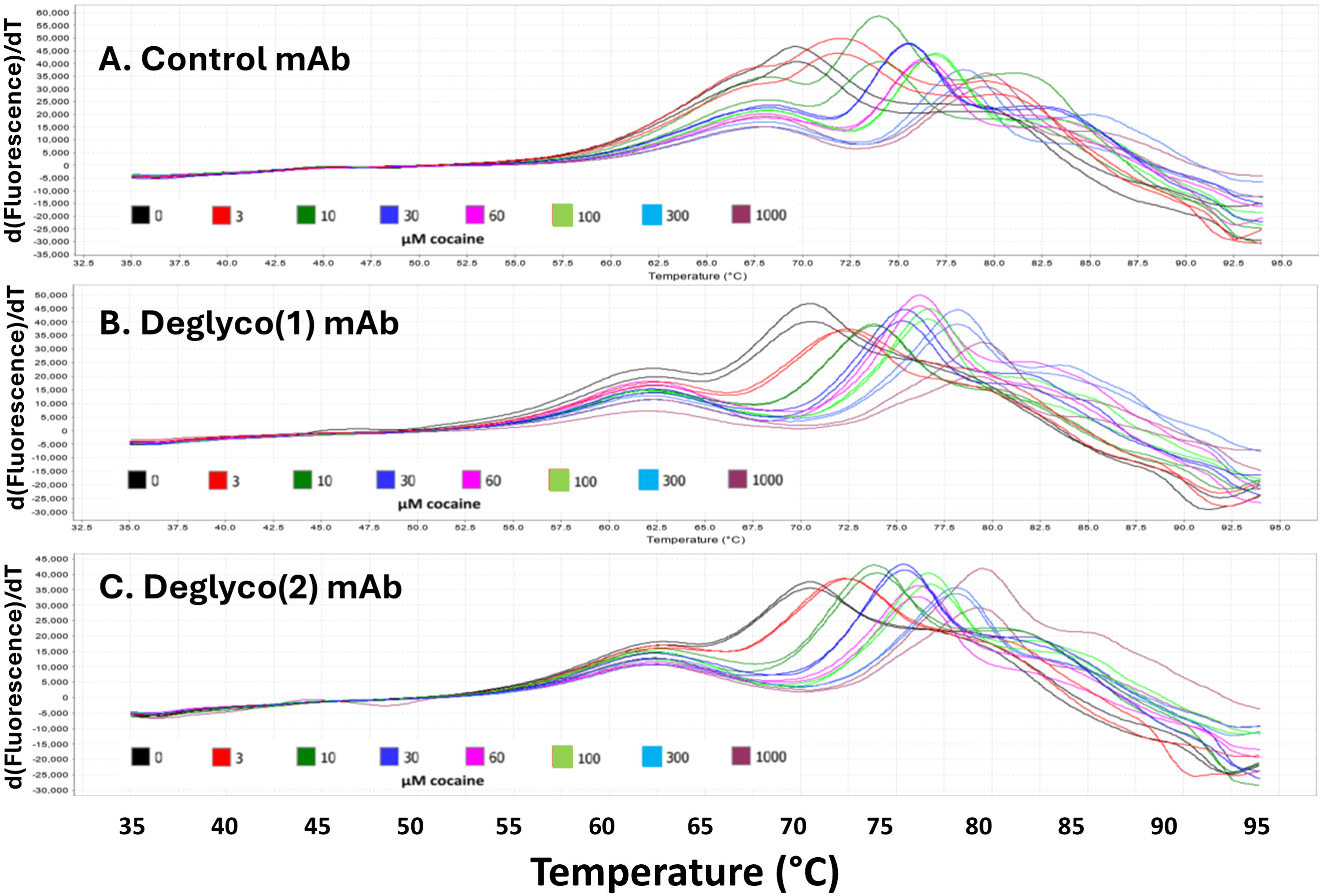
Sypro orange DSF analyses of control and 2 deglycosylated mAb preparations in PBS buffer. All samples contained 2.5 µM mAb, and were analyzed in duplicate, with concentrations of added cocaine varying from 0 to 1000 µM (see color coding in figure). Only the first derivative of the raw fluorescence data are shown, to facilitate comparisons.

**Figure 3.**
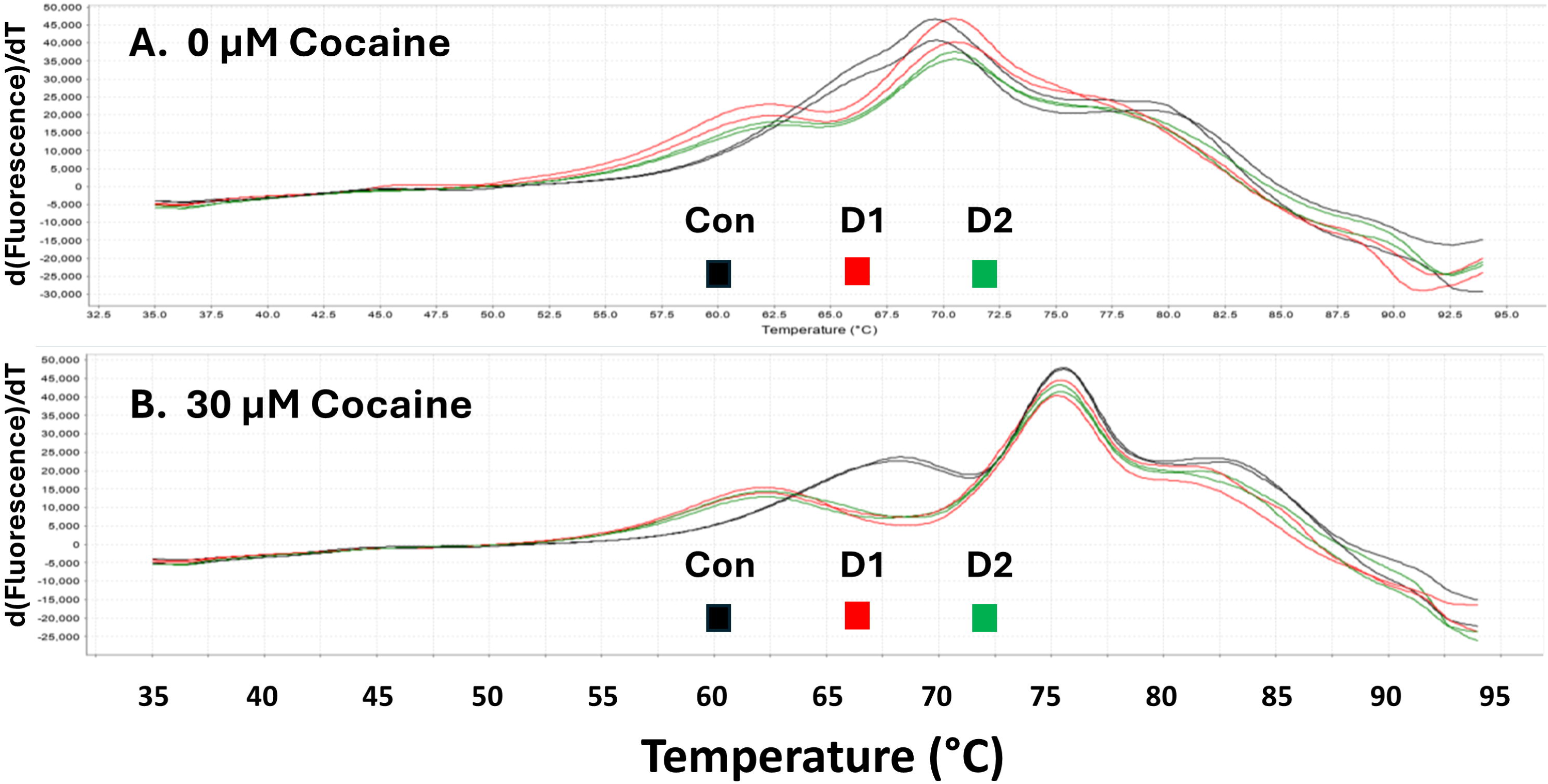
Sypro orange DSF analyses of control and 2 deglycosylated mAb preparations in PBS buffer, comparing samples containing no cocaine with those containing 30 µM cocaine. Data shown is a subset of data shown in Figure 2, to make clear the shift in the lower Tm unfolding transition resulting from deglycosylation.

Figure 4 indicates that results using the DASPMI dye to compare the native and deglycosylated mAb, and the effects of cocaine binding on these samples, are less complex, and indicate little effect of deglycosylation of the Fab portion of the mAb to which DASPMI binds [12]. However, there is a flattening of the peak of the first derivative of the fluorescence signal of the deglycosylated mAb compared to the control, seen most clearly in the absence of cocaine (see Figure 4, compare the black traces in the 3 panels). This derivative shape change caused by deglycosylation is not noticeable in the presence of higher concentrations of cocaine, as seen in Panels B and C, as compared to Panel A, in Figure 4.

**Figure 4.**
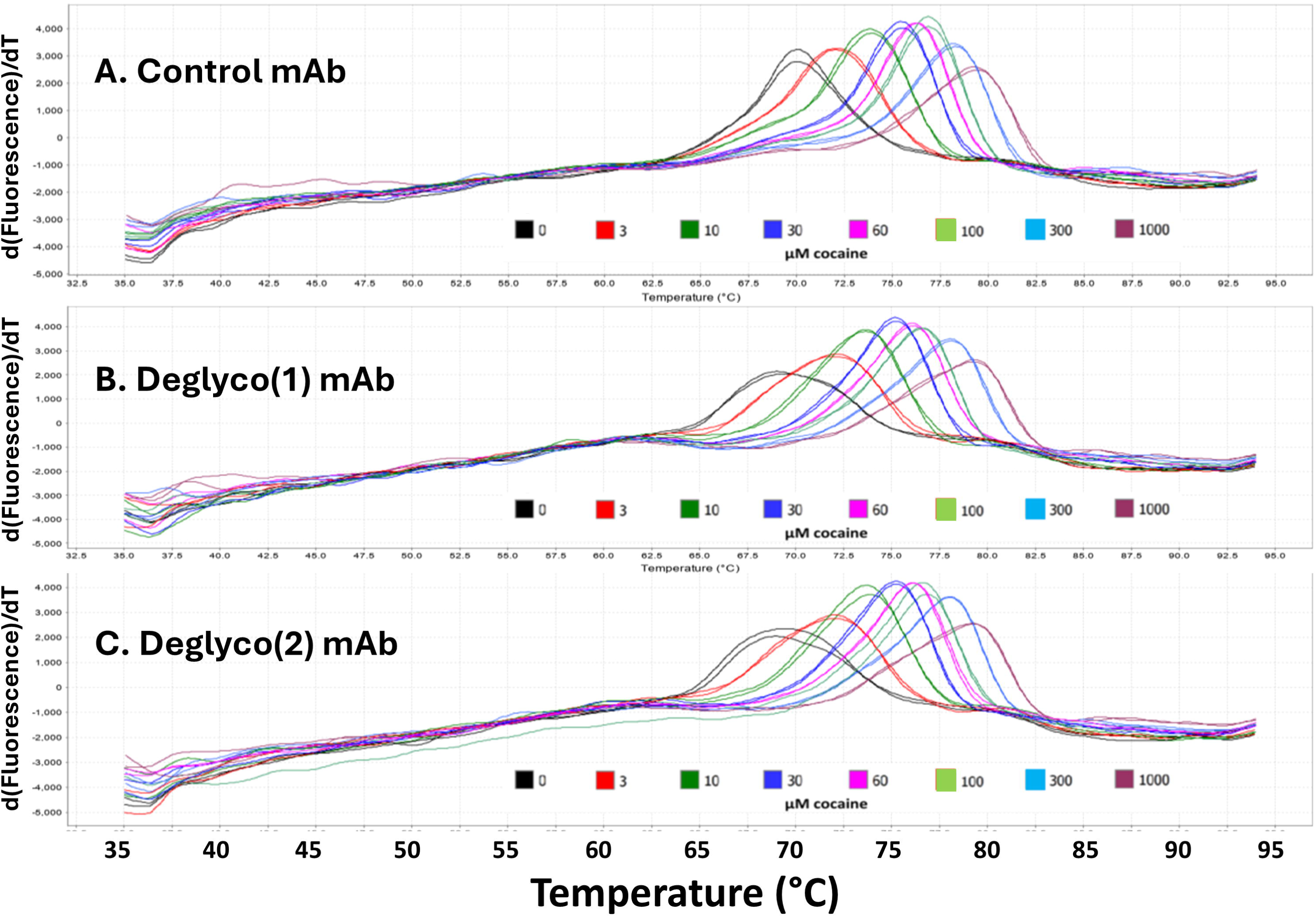
DASPMI DSF analyses of control and 2 deglycosylated mAb preparations in PBS buffer. All samples contained 2.5 µM mAb, and were analyzed in duplicate, with concentrations of added cocaine varying from 3 to 1000 µM (see color coding in figure). Only the first derivative of the raw fluorescence data are shown, to facilitate comparisons.

Control and deglycosylated mAb were labeled with FITC for 30 minutes at 22°C, and the labeled mAb was analyzed by reducing and non-reducing SDS-PAGE, as shown in Figure 5. By absorbance measurements there were virtually no differences in the stoichiometry of FITC labeling (DOL) for these samples, with the DOL being 2.56, 2.44, and 2.63 respectively, for the control C, the D1, and the D2 deglycosylated mAbs. There were also no obvious differences in the relative degree of labeling of the HC or LC parts of the mAb, as well as for the relative degree of FITC labeling for the Fc and Fab fragments following Endo-Lys-C digestion (Figure 5A). In all samples, the majority of the FITC labeling was on the light chain and the Fab fragment, with the Fc fragment being poorly labeled. These same samples were then analyzed using DSF employing the incorporated fluorescein fluorescence, with no extrinsic DSF dye added (Figure 6). As reported previously, the conjugated fluorescein fluorescence decreases with domain thermal unfolding, in contrast to the increase of fluorescence of extrinsic dyes commonly used in DSF, leading to the negative derivative peaks seen in Figure 6 [4]. There were no observable differences in the FITC-mediated DSF between the control and deglycosylated mAbs at all concentrations of cocaine. All samples exhibited the same ligand thermal stabilization of the cocaine binding portion of the Fab fragment to which the DASPMI dye binds. The extremes of cocaine concentrations used in this experiment are shown in Panel B to visually emphasize the lack of glycosylation effects, and the cocaine-induced stabilization of the FITC labeled domain of the mAb.

**Figure 5.**
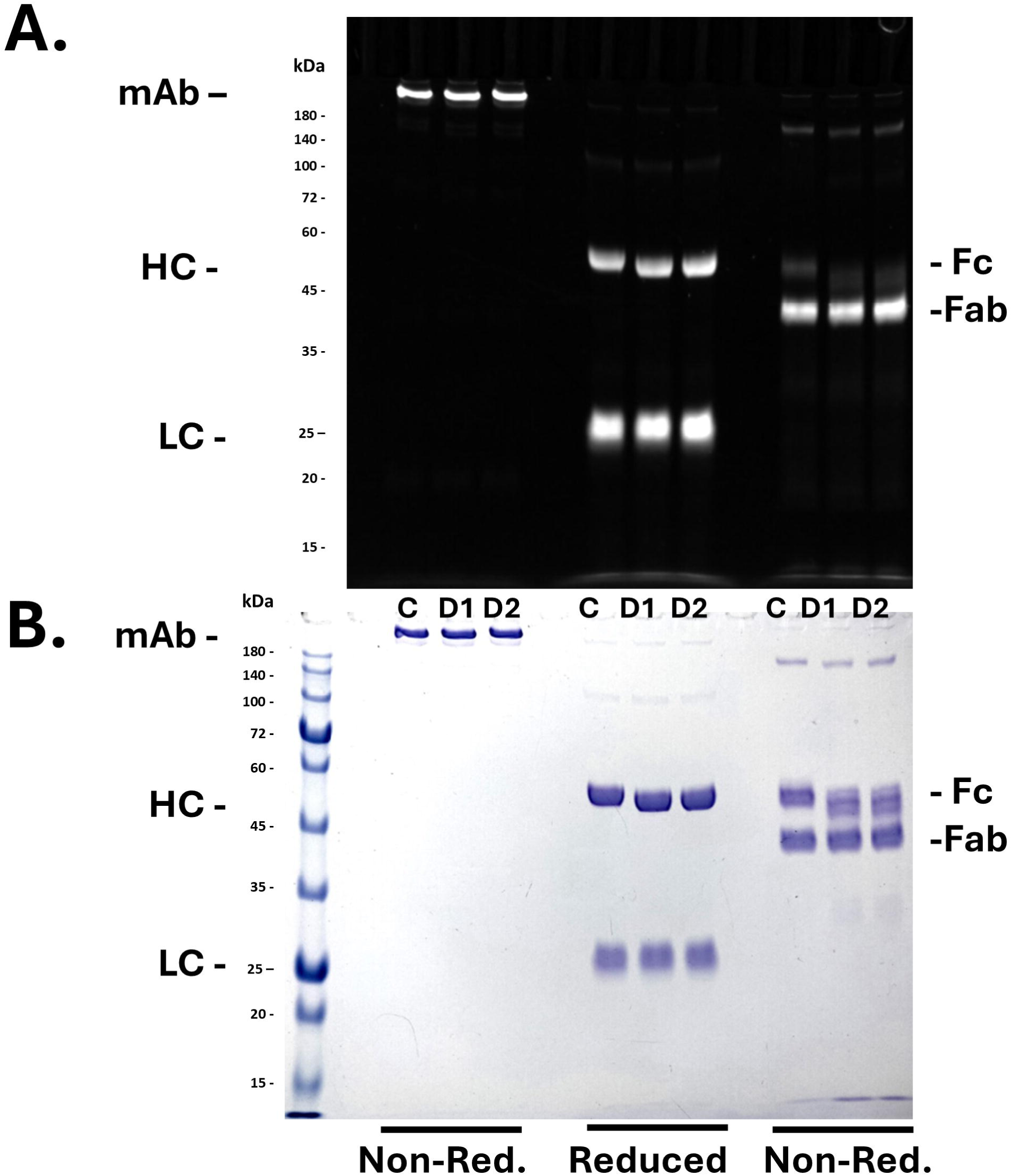
10% SDS-PAGE analysis of FITC labeled control and deglycosylated mAb. 2 µg of each non-reduced intact mAb sample, and 4 µg of reduced intact mAb and non-reduced Endo-Lys-C digests were loaded per gel well. Panel A is the fluorescein fluorescence of the gel bands, while Panel B is the same gel after Coomassie blue staining. The electrophoretic positions of the intact non-reduced mAb, the heavy (HC) and light (LC) chain bands of the reduced mAb, and the Fc and Fab fragments of the mAb digested with Endo-Lys-C (non-reduced) are indicated for the control (C) mAb and two deglycosylated preparations of the mAb (D1 and D2).

**Figure 6.**
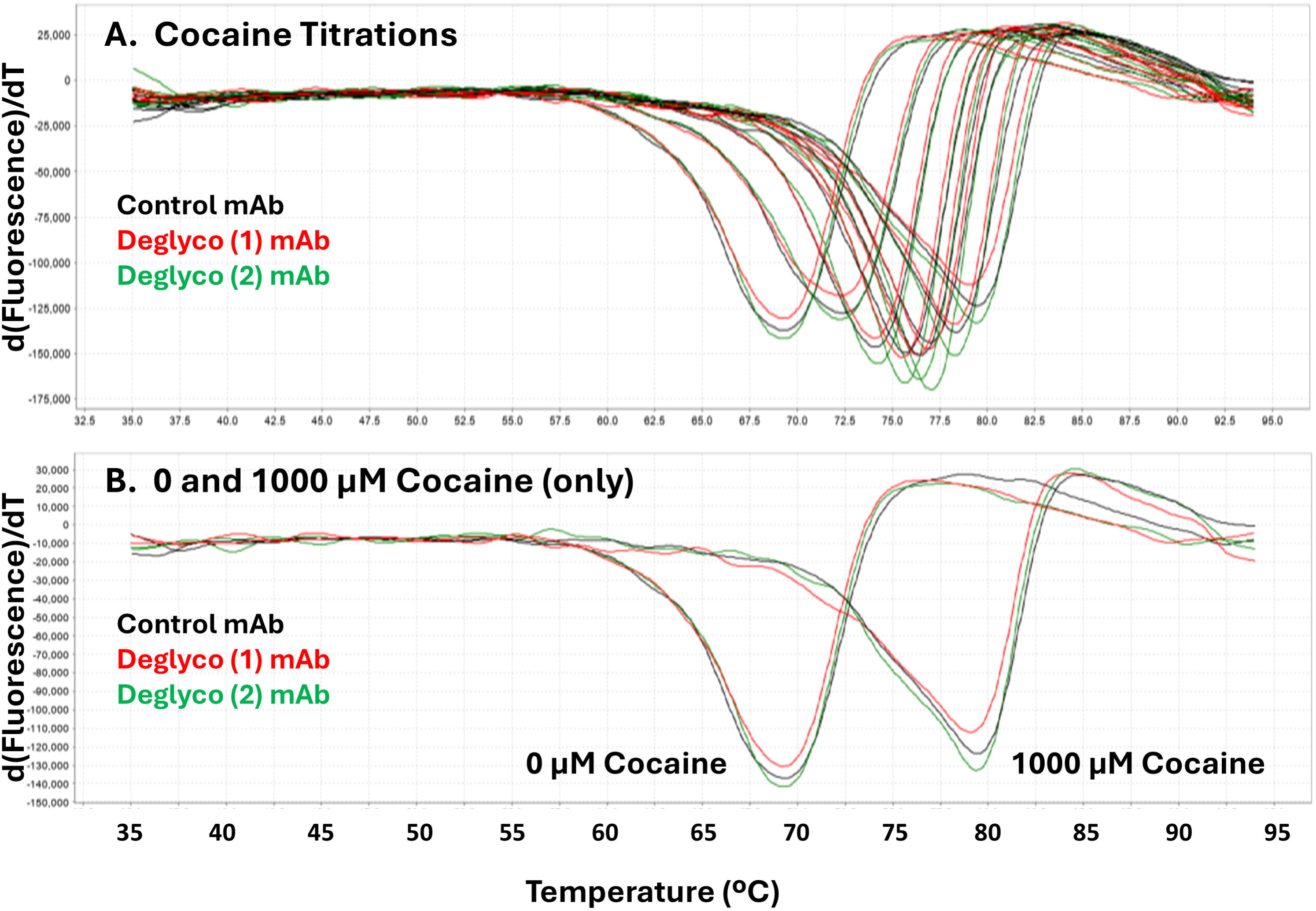
FITC-DSF analyses of control and 2 deglycosylated mAb preparations in PBS buffer. Covalently-incorporated fluorescein fluorescence was monitored as a function of temperature, with all samples containing 2.5 µM mAb, analyzed in duplicate, with concentrations of added cocaine varying from 0 to 1000 µM (Panel A, see color coding for control and deglycosylated mAb samples in figure). The concentrations of cocaine utilized are the same as used in Figures 2 and 4. Only the first derivatives of the raw fluorescence data are shown, to facilitate comparisons. Panel B is a replot of only the 0 and 1000 µM cocaine data from Panel A to facilitate visual comparisons.

Control and deglycosylated mAb were digested with Endo-Lys-C, after performing a time course of digestion to establish optimum conditions for generation of Fc and Fab fragments (preliminary time course data not shown). The Fc fragment was then purified from the digest by cation exchange chromatography, utilizing the differences between its pI value of 7.0, compared to the pI of the Fab fragment (8.57) and the intact mAb (8.37), see [14]. The resultant purified Fc fragments were analyzed by reducing and non-reducing SDS-PAGE (Figure 7). Again, under reducing conditions, the deglycosylation was demonstrated to be complete, and as expected. However, Endo-Lys-C digestion of the deglycosylated Fc led to the production of Fc sub-fragments (labeled Fc (1) and Fc (2) in the Figure). In addition, there appears to be some heterogeneity in the deglycosylated Fc fragment, since multiple bands are observed under non-reducing conditions for the deglycosylated Fc, unlike the glycosylated Fc. The asterisks (*) in the figure mark the migration position of the Fab fragment, indicating that there remains a small amount of Fab contamination, despite two rounds of Fc ion exchange purification. The 2 lower molecular weight bands observed in the deglycosylated Fc sample are almost certainly generated by an Endo-Lys-C proteolysis site that is exposed by deglycosylation, since the gel determined molecular weights of the two sub-fragments observed add up to the gel molecular weight of the deglycosylated Fc (i.e., 27.8 kDa = 15.6 kDa + 12.2 kDa (see molecular weight calibration and determination data shown in Figure 8, which are derived from the gel bands shown in Figure 7)). Control and deglycosylated Fc fragments were analyzed using both Sypro orange and DASPMI dyes (Figure 8). The presence of cocaine did not affect these Fc data, as expected (cocaine data not shown). The Sypro orange dye did detect multiple Fc thermal unfolding transitions, the most evident one having a Tm of about 68.5°C in the glycosylated Fc (Figure 9A, black traces), which transition is not observed in the deglycosylated Fc. Instead, in the deglycosylated Fc there are a couple of very broad, lower Tm thermal transitions centered around 48°C and 58°C (red traces in Figure 9A). The DASPMI dye detected no thermal transitions in either the control or the deglycosylated Fc (Figure 9B), which was expected, given previous results indicating that the DASPMI dye does not bind to the Fc fragment of this anti-cocaine mAb [12].

**Figure 7.**
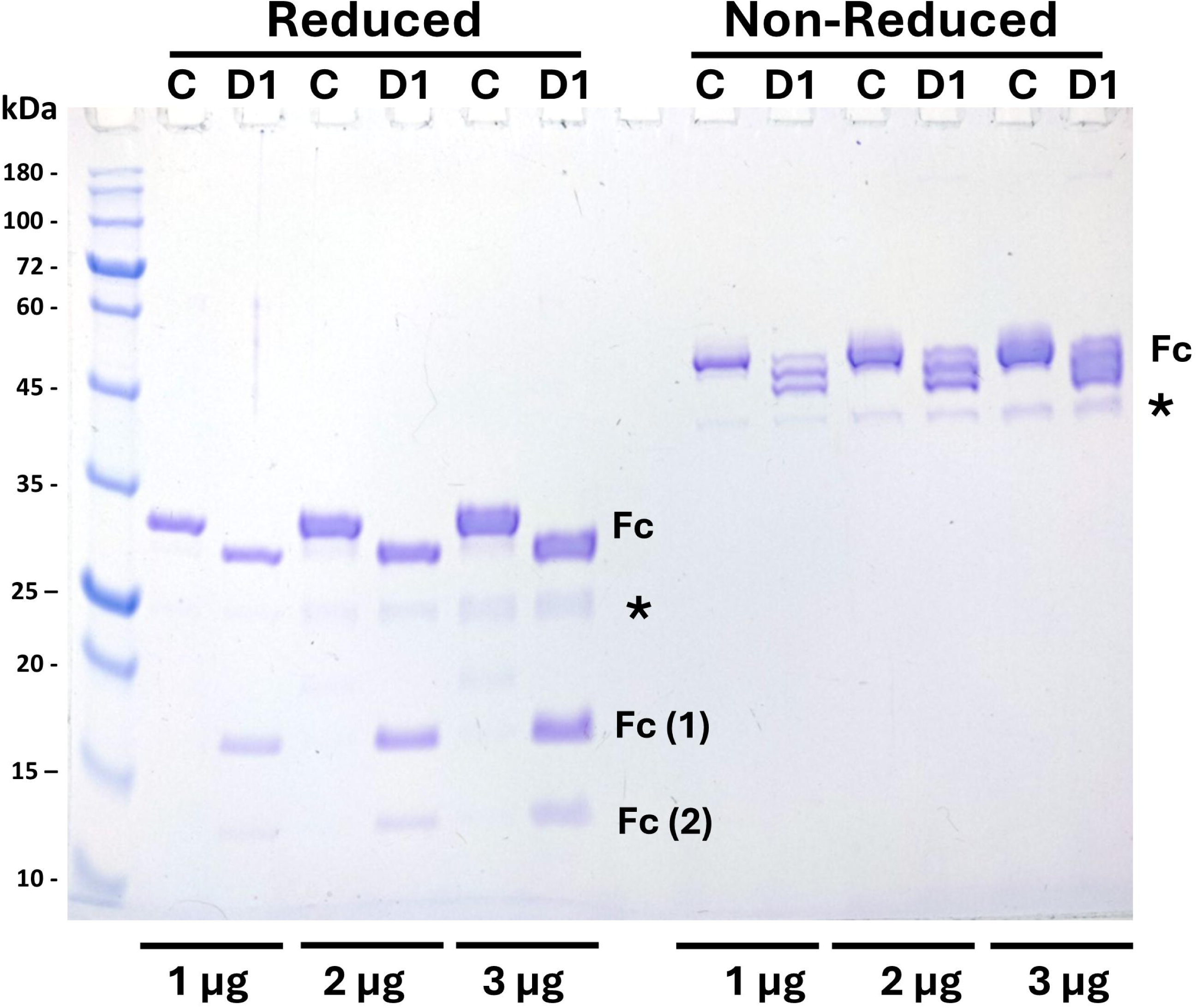
12% SDS-PAGE analysis of control and deglycosylated purified Fc fragments under both reduced and non-reduced conditions. 1, 2, or 3 µg Fc fragments were loaded on the gel as described in methods for control (C) mAb or deglycosylated mAb (D1, deglycosylated with NEB PNGase F) after digestion with Endo-Lys-C, and purification. Reduced and non-reduced Fc bands are annotated, as well as the 2 Fc sub-fragments seen only in the deglycosylated sample. A very small amount of contaminating Fab fragment is indicated by an “*”. Note the shift to lower molecular weight for the reduced Fc fragment, and the multiple bands observed for the deglycosylated, non-reduced Fc fragment in the deglycosylated sample.

**Figure 8.**
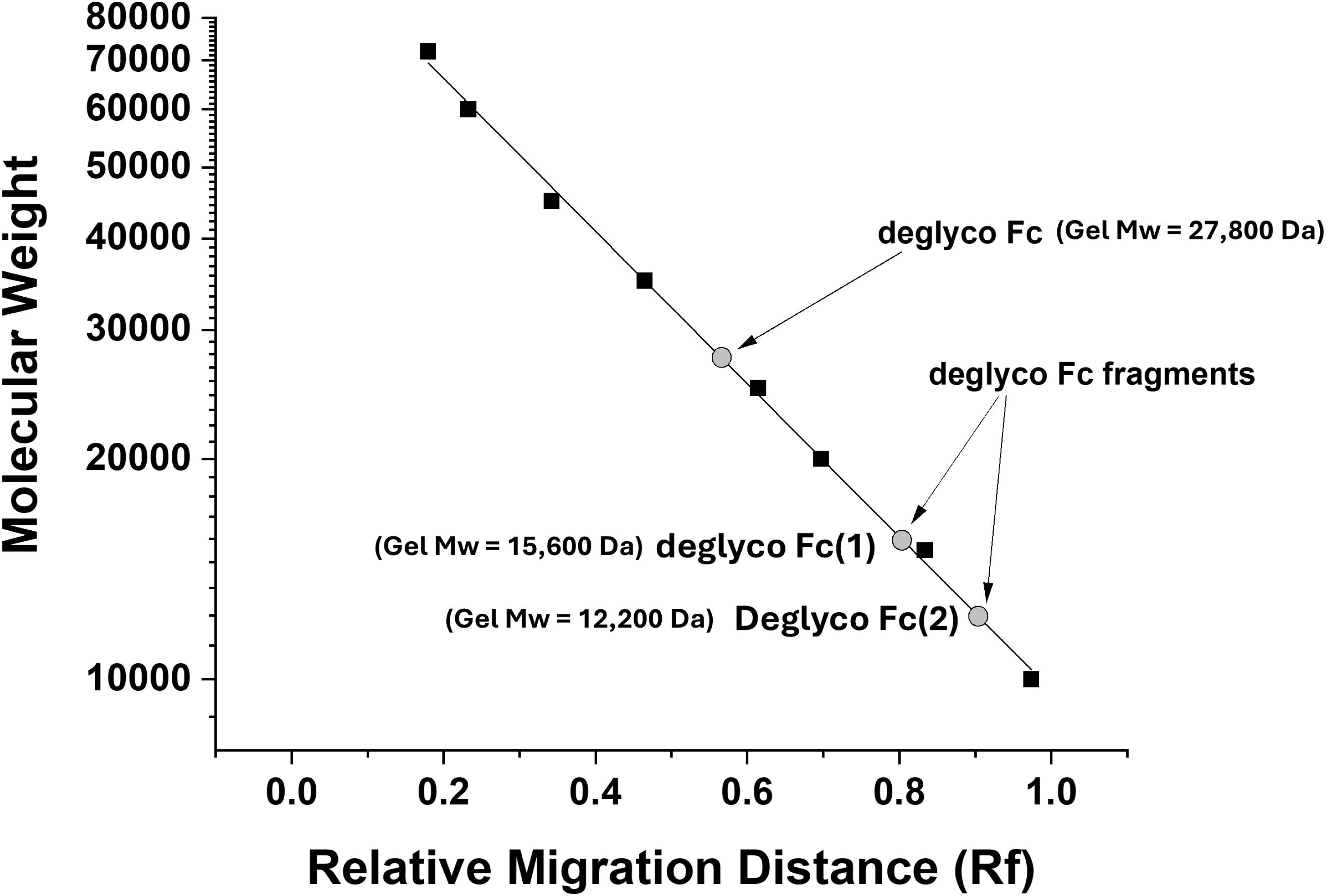
Sizing of the deglycosylated Fc band and Fc sub-fragment bands seen in Figure 7. Molecular weight standard protein relative migration distances (Rf) on the gel are plotted versus the log of the molecular weight of these standard proteins. The apparent sizes of the deglycosylated Fc band, as well as the two Fc sub-fragments of the deglycosylated Fc were calculated from the straight line fit to standard proteins less than 100 kDa on the 12% SDS-PAGE, and the Fc and sub-fragment molecular weights are indicated in the figure.

**Figure 9.**
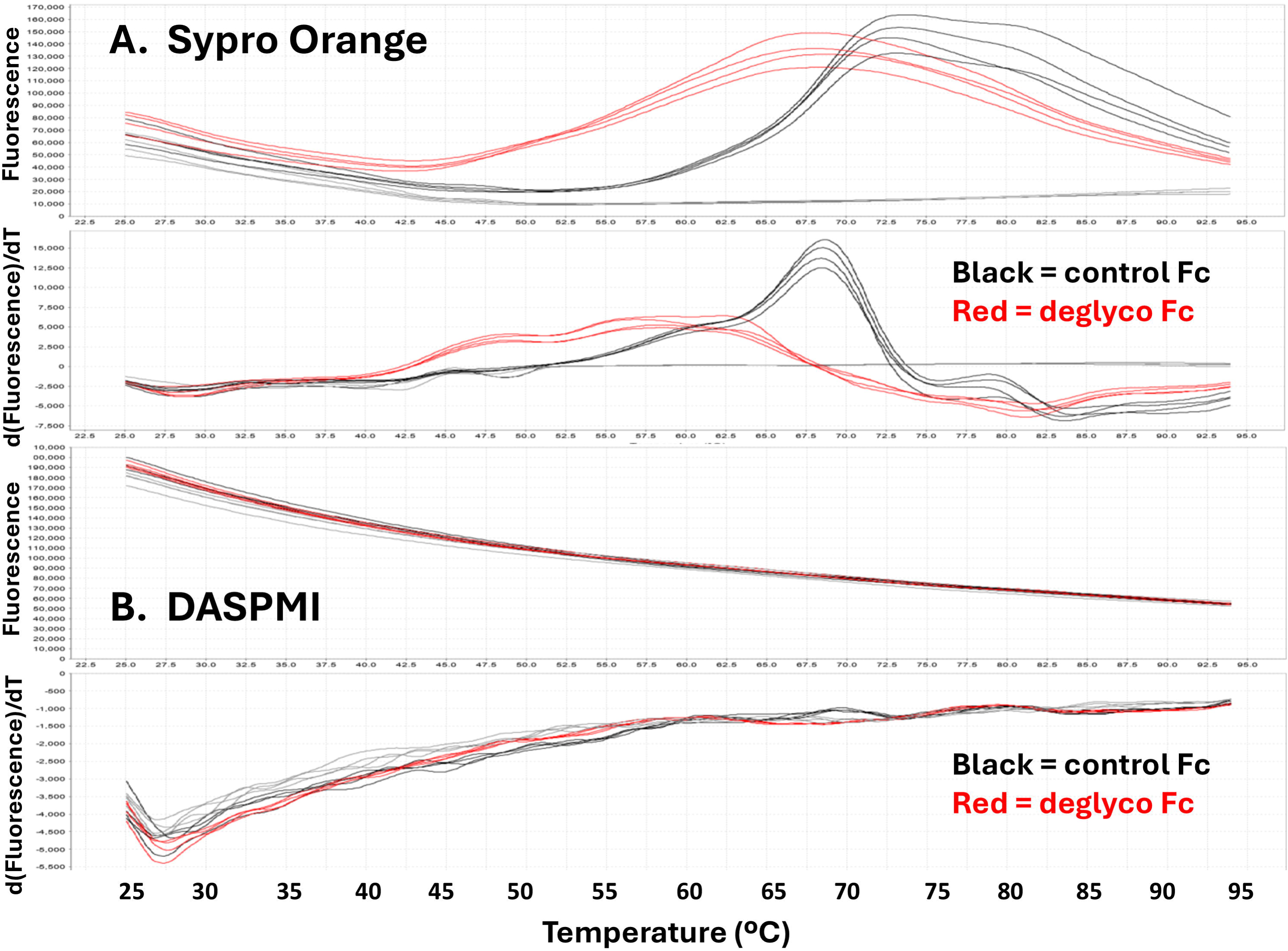
Sypro orange and DASPMI extrinsic dye DSF analyses of control and deglycosylated Fc in PBS buffer. Control and deglycosylated purified Fc fragments are compared in the figure. Panel A is the Sypro orange raw fluorescence (top) and the first derivative of that fluorescence (bottom) as a function of temperature, while Panel B is the DASPMI dye raw fluorescence (top) and the first derivative of that fluorescence (bottom) as a function of temperature. All samples were analyzed in quadruplicate wells

The control and deglycosylated Fc fragments were labeled with FITC under two different reaction conditions, as described in the methods section. These labeled proteins were analyzed by SDS-PAGE under both reduced and non-reduced conditions (Figure 10). The DOL (mol FITC/mol Fc fragment) for the lesser degree of FITC labeling for 1 hour in MOPS buffer 7.4 was 0.11 for control (C1) and 0.37 for deglycosylated Fc (D1), while for the greater degree of FITC labeling for 2 hours in borate buffer pH 8.0 was 0.25 for control (C2) and 0.65 for deglycosylated Fc (D2), as determined by absorbance measurements. Thus, in both sets of FITC labeled samples, it is obvious that the amount of fluorescein labeling was greatly increased in the deglycosylated Fc compared to the control Fc, and that most of this increase in FITC labeling is due to labeling of the larger, Fc(1), deglycosylated Fc sub-fragment (Figure 10A). These 4 samples were also analyzed using FITC-mediated DSF, as shown in Figure 11. The various levels of total FITC labeling of the samples are reflected in the different levels of starting (25°C) fluorescence in the raw fluorescence traces shown in the top graph. As is evident, there is a fluorescein-detected thermal transition in each of the deglycosylated Fc samples at about 79°C (red and pink traces in the bottom, first derivative graph), which are almost absent in the control, glycosylated FITC labeled Fc (black and gray traces).

**Figure 10.**
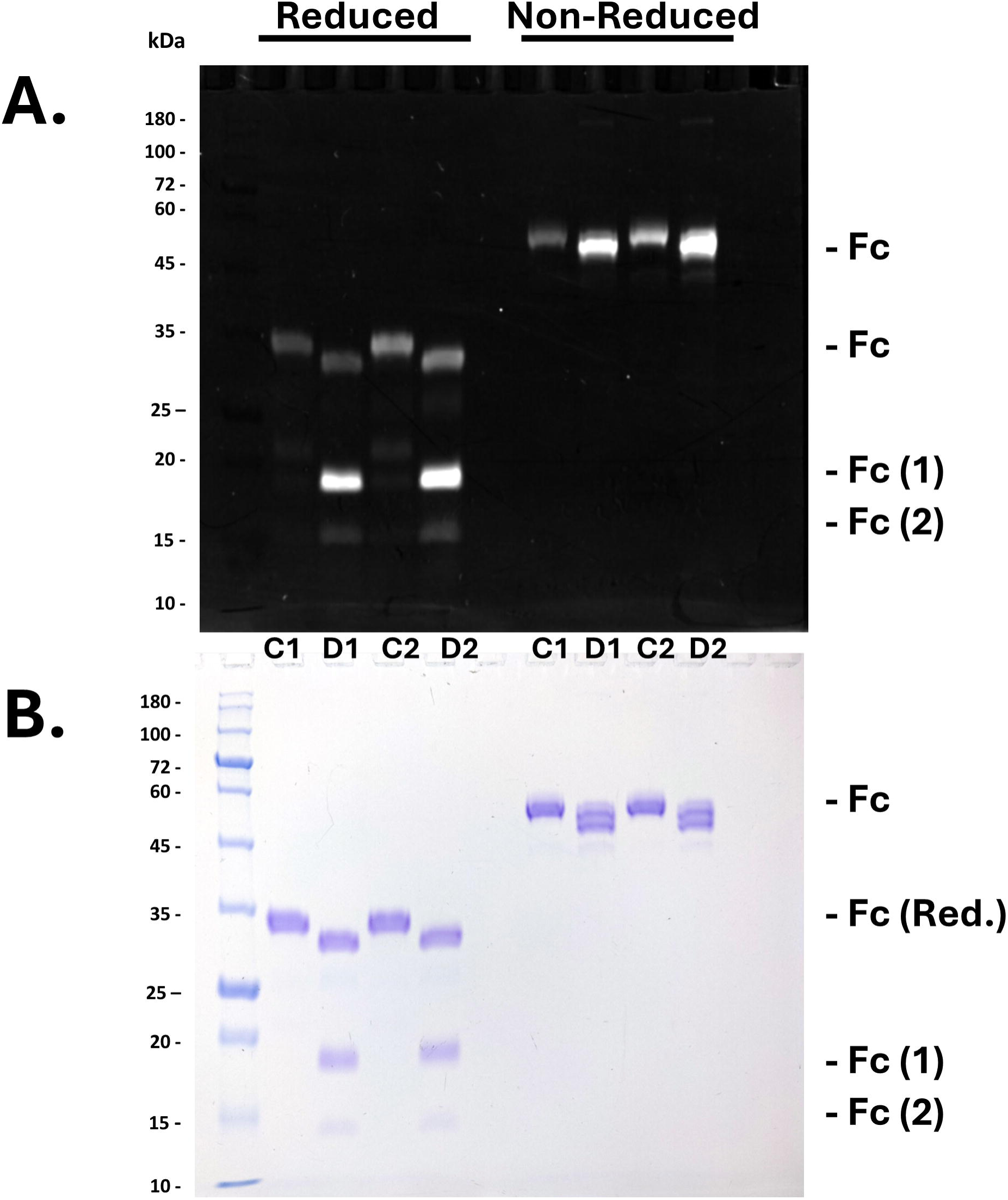
Reducing and non-reducing 12% SDS-PAGE analysis of FITC labeled control and deglycosylated purified Fc fragments. 2 µg of each sample was loaded per gel well. Panel A is the fluorescein fluorescence of the gel protein bands, while Panel B is the same gel after Coomassie blue staining. The electrophoretic positions of the Fc fragments, as well as the two Fc fragments observed only in the deglycosylated Fc sample are annotated. C1 and D1 are the control and deglycosylated Fc labeled to a lower level with FITC in MOPS buffer, while samples C2 and D2 are the control and deglycosylated Fc labeled to a higher level with FITC in borate buffer.

**Figure 11.**
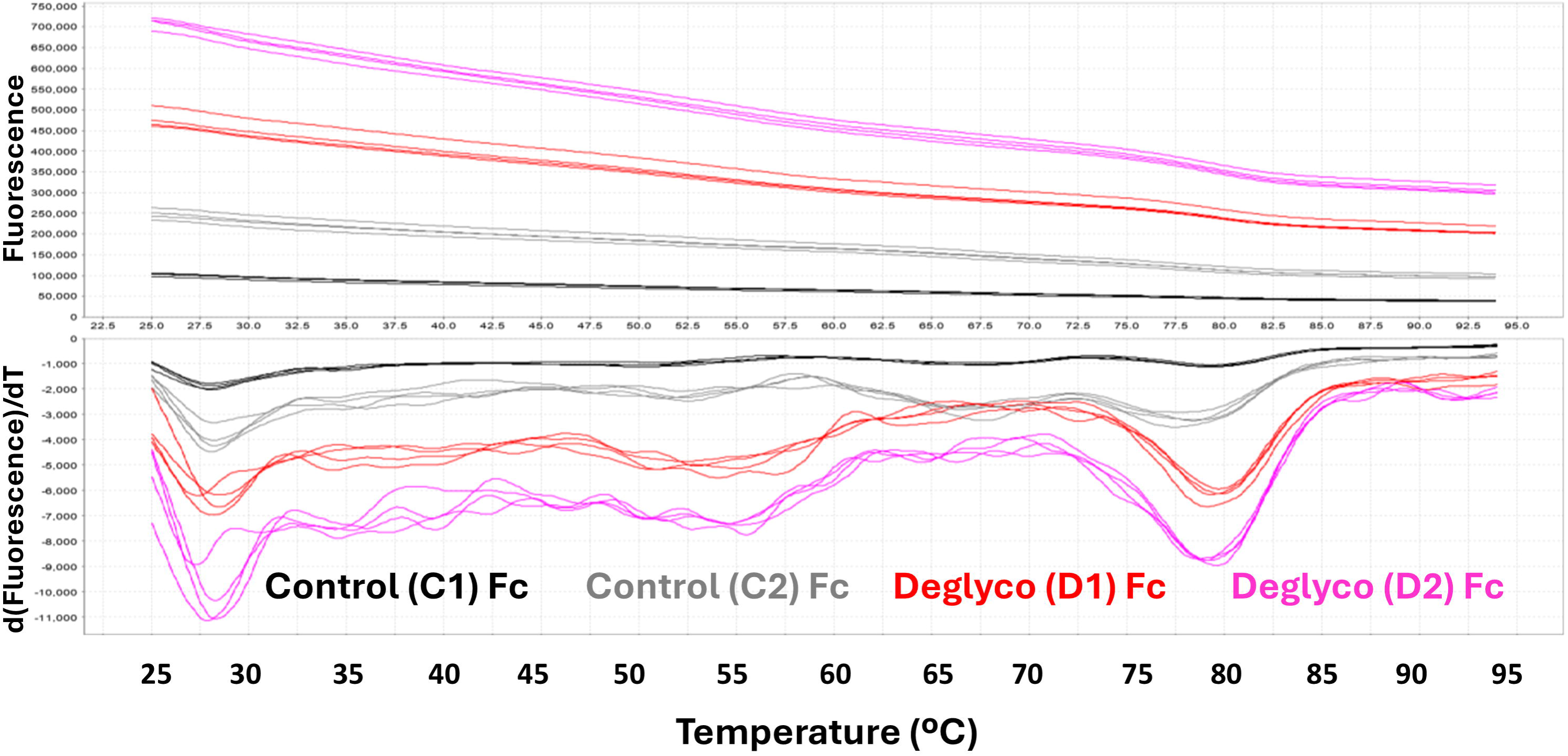
FITC DSF analyses of control and deglycosylated Fc preparations in PBS buffer. FITC raw fluorescence (top) and the first derivative of that fluorescence (bottom) is shown as a function of temperature for two sets of Fc samples (control and deglycosylated) labeled with under different conditions, Control (C1) Fc and Deglyco (D1) Fc were FITC labeled to a lesser level in MOPS buffer, whereas Control (C2) Fc and Deglyco (D2) Fc were FITC labeled to a higher level in borate buffer. Samples were analyzed in quadruplicate wells.

## 4. Discussion

The objective of this study was to characterize the effect of deglycosylation on the properties and thermal stability of the fragmented and intact h2E2 anti-cocaine mAb under development for the treatment of cocaine use disorders, using several variations of differential scanning fluorimetry (DSF). This characterization is vital, since antibody glycosylation has been shown to be important for conformational stability [11], effector functions [15], and pharmacokinetics and half-lives of therapeutic mAbs [6]. Also, the unfolding of the (glycosylated) CH2 domain of the Fc region has been implicated as the domain whose unfolding triggers irreversible mAb aggregation [16], which is a very important issue for the stability, storage, and usage of therapeutic mAbs. Differential scanning calorimetry (DSC) as well as DSF employing the Sypro orange dye are commonly used to assess antibody protein domain stabilities, and the effects of mutation, chemical modification, and drug conjugation on those domain stabilities, as well as the effects of solvent excipients used in the formulation of therapeutic mAbs [17]. Sypro orange DSF has been employed to predict thermal aggregation of proteins [18], and in general, DSC and DSF results for antibodies indicate that the order of stability of antibody domains, from least stable (lowest Tm) to most stable (highest Tm) progresses from the Fc CH2 domain (which is glycosylated in all IgG1 mAbs) to the Fab domain to the Fc CH3 domain [19], although Fab domain thermal stabilities of mAbs can vary widely. Thus, the stability and glycosylation of the CH2 domain of the Fc fragment is especially important for the thermal stability, aggregation tendency, and usage of many experimental and therapeutic antibodies.

DSF typically uses extrinsic dyes, which change their fluorescence characteristics or emission yields when bound to hydrophobic patches of protein structure that are exposed by thermal unfolding of the protein or domains of the protein (for reviews and applications of DSF and properties of many of the dyes used in DSF, see [20–22]). The most commonly used type of dyes for DSF are solvatochromatic dyes, whose fluorescence emission increase dramatically when bound to a hydrophobic patch on a protein, which becomes accessible during thermal unfolding. Most DSF studies employ Sypro orange as the solvatochromatic dye. As expected, the solvatochromatic Sypro orange dye binds to several domains of the h2E2 anti-cocaine mAb, including the Fc and Fab domains, resulting in multiple, overlapping thermal unfolding transitions being detected in Sypro orange DSF for this mAb [12]. We have championed the use of the rotor DASPMI DSF dye, which is characterized by its fluorescence increasing when its rotational freedom is restricted by binding to regions of the protein that become accessible upon protein domain unfolding, and does not bind to the Fc fragment of the anti-cocaine mAb. DSF analyses utilizing the DASPMI dye have been used to quantitate the affinity of cocaine and cocaine metabolites to the h2E2 anti-cocaine mAb [12]. In addition, limited fluorescent labeling with FITC was recently shown to be useful for DSF analyses in the absence of added extrinsic dyes to detect cocaine-induced stabilization of the antigen binding domain [4].

To investigate the importance and effects of glycosylation on the anti-cocaine mAb, the glycan chains on each heavy chain were quantitatively removed using PNGase-F (Figure 1), and the resultant deglycosylated mAb was analyzed and compared to the control mAb using Sypro orange DSF, revealing several overlapping thermal transitions, one of which, occurring at the lowest temperature, was shifted to an even lower temperature by deglycosylation. This deglycosylation destabilization could be better detected in the presence of cocaine, which stabilized the Fab domain and shifted its thermal transition to higher temperatures (Figure 2). This phenomenon was reproduced using 2 different commercially available PNGase-F deglycosylation enzymes (Figure 2B and 2C). The degree of thermal destabilization of the lower temperature denaturing domain is clearly seen in Figure 3, with the Tm decreasing from about 68°C to about 62°C after deglycosylation, indicating a large, reproducible decrease in the thermal stability of that CH2 mAb domain.

In contrast, deglycosylation had minimal effect on the part of the Fab fragment that binds cocaine, as demonstrated by the DASPMI DSF data shown in Figure 4. However, it should be noted that in the absence of cocaine, the DASPMI DSF derivative traces are reproducibly broader after deglycosylation (compare the black traces in the figure). The reason for this is unclear, but this effect decreases and then disappears with increasing concentrations of cocaine stabilizing this domain, as seen in the figure.

Fluorescent labeling with FITC is a widely used chemical modification of proteins and antibodies, so the effects of deglycosylation on the labeling of the mAb (and its Fc and Fab fragments) with FITC was examined. Consistent with earlier results [4], there was more FITC labeling on the light chain than the heavy chain of this mAb, and also more labeling on the Fab fragment than the Fc fragment (Figure 5). However, neither the total FITC level incorporated, nor the distribution between these mAb chains or domains, was different after deglycosylation of the mAb (Figure 5). This was also true when increasing FITC labeling by increasing the time of labeling from 30 minutes to 2 hours. Thus, after 2 hours of FITC labeling at 22°C, there were also virtually no differences in the stoichiometry of FITC labeling (DOL) for these samples, with the DOL being 4.56, 4.49, and 4.65 respectively, for the control C, the D1, and the D2 deglycosylated mAbs. In addition, gel analyses of the relative FITC 2-hour labeling of HC vs LC and Fab vs Fc fragment labeling also displayed no differences between the native and deglycosylated mAb (2 hour FITC labeled gel data not shown).

FITC-mediated DSF analysis and the effect of cocaine on FITC DSF results were also not different for the control versus the deglycosylated mAb (Figure 6). In all cases, the temperature at which the decrease in fluorescein fluorescence covalently bound to the mAb occurs is shown to increase with increasing cocaine concentrations, indicating that FITC labeled mAb can be used to qualitatively assess the binding of cocaine and its metabolites. This is due to most of the fluorescein being covalently bonded to the Fab portion of the antibody (Figure 5A), which binds cocaine, but is far removed from the glycosylation sites on the heavy chain CH2 domain, and thus FITC labeling under these conditions is not useful to assess effects of deglycosylation in the Fc CH2 domain of the intact mAb.

Therefore, to investigate the effects of deglycosylation more directly, the control and deglycosylated mAb were cleaved into their respective Fc and Fab fragments using Endo-Lys-C, and the Fc fragments were purified and analyzed. One unanticipated result is the occurrence of an additional site of limited Endo-Lys-C cleavage on the deglycosylated Fc, forming two deglycosylated Fc sub-fragments, designated Fc (1) and Fc (2) (Figures 7 and 8). By gel-determined molecular weight, Fc (1) fragment comprises 56.1% of the deglycosylated Fc molecular weight (15,600 Da/27,800 Da). Applying this percent molecular mass to the calculated theoretical Endo-Lys-C fragment amino acid sequence of the heavy chain, assuming a C-terminal lysine residue at the secondary site of proteolytic cleavage, the best Fc HC amino acid sequence fragment size “fit’ to explain this Fc (1) sub-fragment sizing data is : C**K**VSN**K**ALPAPIEKTISKAKGQPREPQVYTLPPSREEMTKNQVSLTCLVKGFYPSDIAVE WESNGQPENNYKTTPPVLDSDGSFFLYSKLTVDKSRWQQGNVFSCSVMHEALHNHYTQKSLSLSPGK, which is also 56.1% of the total HC Fc theoretical molecular weight of the Endo-Lys-C amino acid sequence (14.22 kDa/25.34 kDa, masses calculated based on the amino acid sequences). This means that the most likely secondary Endo-Lys-C cleavage point would occur at the last (bolded and underlined) lysine (K) residue in the following sequence from the heavy chain (the consensus conserved N-linked glycosylation site sequence is underlined): …QY<u>NST</u>YRVVSVLTVLHQDWLNG**K**EY**<u>K</u>** (this sequence, which would be part of the Fc(2) sub-fragment sequence, immediately precedes the Fc(1) sequence shown above). However, it should be noted that there are several additional lysine residues located near this putative site (3 lysine residues within 5 residues of the putative lysine cleavage site - see K residues which are bolded in the above sequences). This clustering of lysine residues would likely lead to secondary cleavage at several different lysine residues in this region, which may explain the observed SDS-PAGE heterogeneity of the non-reduced deglycosylated Fc fragment seen in Figure 7, especially since there is a cysteine residue involved in a disulfide bond which is in close proximity to these putative lysine cleavage sites. Cleavage on one side of that disulfide versus cleavage on the other side of that disulfide could lead to differential SDS-denatured protein shapes, slightly changing its electrophoretic migration rate on non-reducing SDS-PAGE, explaining the heterogeneity in the non-reduced deglycosylated Fc band shown in Figure 7.

As anticipated, the DASPMI dye did not bind to, or report on the stability of, the purified Fc fragment (Figure 9B). However, the Sypro orange DSF revealed a major Fc transition with a Tm of 68.5°C, which is not present after deglycosylation, presumably due to destabilization of the Fc CH2 domain. Instead, the deglycosylated Fc exhibits extremely broad, lower Tm transitions at approximately 48°C and 58°C (red traces in Figure 9A). This indicates a very large destabilization of the isolated Fc CH2 domain by deglycosylation.

The secondary site of proteolysis on the Fc fragment that was exposed by deglycosylation led to a large increase in FITC labeling of the deglycosylated Fc under two labeling conditions, due to labeling of the larger, Fc (1) sub-fragment (Figure 10A). This increase in labeling is likely due to lysine labeling at or near the secondary cleavage site lysine(s). Regardless, this increased labeling resulted in a FITC-mediated DSF thermal transition being detected in the deglycosylated Fc but not in the glycosylated Fc, with a Tm of 79.0°C (Figure 11). This thermal transition is due to the unfolding of the thermally stable CH3 mAb domain, which is contained in the amino acid sequence of the Fc (1) fragment. Thus, FITC labeling DSF enables much better observation of this transition than that seen with the Sypro orange dye, where this transition at about 79°C is seen only as an unresolved, overlapping bump in the DSF traces shown in Figure 3A. The CH2 domain thermal unfolding transition for the control Fc detected by Sypro orange (the black traces in Figure 9A, with a Tm of about 68.5°C) is not detected by the FITC-mediated DSF shown in Figure 11, since there is very little FITC labeling of the control Fc fragment or of the deglycosylated Fc (2) sub-fragment (see Figure 10A) that includes the amino acid sequence comprising the CH2 domain of the Fc fragment.

## 5. Conclusions

Analyses of the anti-cocaine mAb by DSF utilizing 2 distinct external dyes, Sypro orange and DASPMI, and also the fluorescently labeled protein following limited FITC labeling of mAb lysine residues, were performed with and without deglycosylation. The effects of binding of the antigen, cocaine, on these DSF thermal transitions were monitored and utilized to differentiate and better identify some of the multiple mAb and Fc domain unfolding transitions, as compared to results obtained with the Sypro orange dye typically used in protein DSF experiments. Deglycosylation resulted in a major destabilization of the Fc fragment, as well as the exposure of a secondary site of limited Endo-Lys-C cleavage during proteolytic production of the Fc fragment from the deglycosylated mAb, which allowed FITC-mediated DSF analysis of the deglycosylated Fc fragment. Some of the methods used and results obtained should be generalizable to many commonly utilized IgG1 mAbs, since the IgG1 h2E2 anti-cocaine mAb used in this study shares the same conserved CH2 glycosylation site within a fully human heavy chain Fc sequence with other human and humanized IgG1 mAbs used in the laboratory and the clinic.

## Abbreviations

mAb: monoclonal antibody
h2E2: humanized anti-cocaine monoclonal antibody
DSF: differential scanning fluorimetry
DASPMI: (4-(4-(dimethylamino)styryl)-N-methylpyridinium iodide)
Sypro: Sypro Orange dye
PNGase-F: peptide N-glycosidase-F
D or deglyco: deglycosylated (with PNGase-F) mAb
D1: PNGase-F mAb preparation #1
D2: PNGase-F mAb preparation #2
Con or C: Control
SDS-PAGE: sodium dodecyl sulfate polyacrylamide gel electrophoresis
MOPS: 3-(N-morpholino)propanesulfonic acid
PBS: phosphate buffered saline, pH=7.4
DTT: dithiothreitol
FITC: fluorescein isothiocyanate
DOL: degree of labeling, mol label/mol protein
DMSO: dimethyl sulfoxide
Endo-Lys-C: Endoproteinase Lys-C
Fab: fragment antigen-binding
Fc: fragment crystallizable
LC: antibody light chain
HC: antibody heavy chain
Tm: protein domain melting temperature
TmD: first derivative-derived melting temperature
TmB: Boltzmann fitting-derived melting temperature
Kd: binding dissociation constant

## Acknowledgments

I acknowledge the National Institutes of Health National Institute on Drug Abuse Grant U01DA050330, awarded to Dr. Andrew Norman in this Department (Department of Pharmacology, Physiology, and Neurobiology at the University of Cincinnati College of Medicine), which previously funded the production of the recombinant humanized h2E2 anti-cocaine mAb protein by Catalent PharmaSolutions, Inc. (Madison, WI), and also the purchase of some reagents used in this study. Thanks also to Dr. Guochang Fan (also in the Department of Pharmacology, Physiology, and Neurobiology at the University of Cincinnati College of Medicine) for the use of the StepOne RT PCR instrument used for DSF analyses.

